## Supplementary material for "MorphoCellSorter: An Andrews plot-based sorting approach to rank microglia according to their morphological features": SuppData.pdf

**Supp. figure 1**

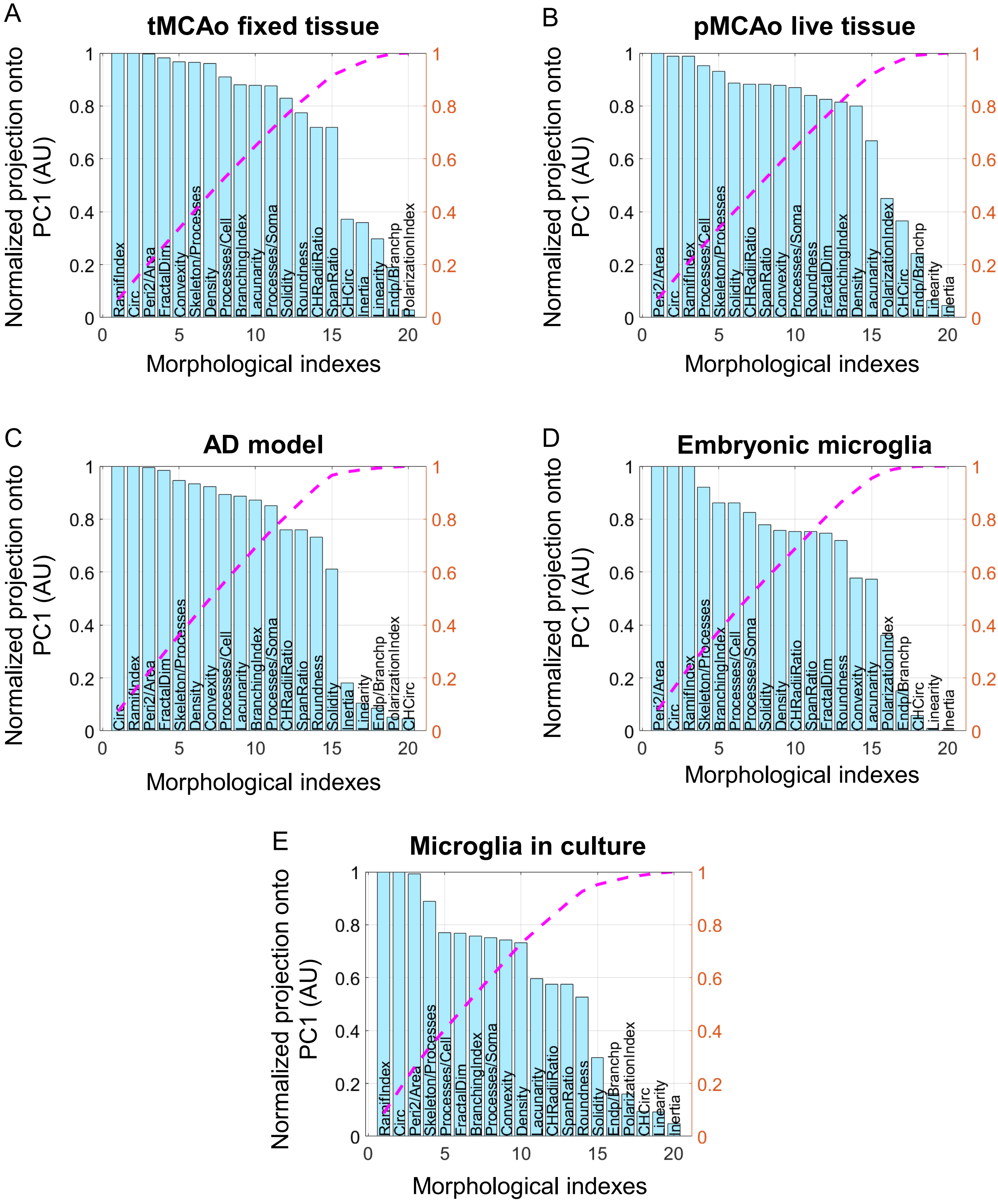

**Supp. table 1**

| Th | Sum $R_S$ experts 1 and 2 | | | | | Threshold ranks | | | | | Sum ranks |
|---|---|---|---|---|---|---|---|---|---|---|---|
| | tMCAo | pMCAo | AD | EM | MC | tMCAo | pMCAo | AD | EM | MC | |
| **0.2** | 1,92325 | 1,88991 | 1,88553 | 1,79939 | 1,6033 | 4 | 1 | 9 | 8 | 7 | 29 |
| **0.3** | 1,91802 | 1,88962 | 1,89456 | 1,79939 | 1,6033 | 7 | 2 | 4 | 8 | 7 | 28 |
| **0.4** | 1,92867 | 1,87351 | 1,89725 | 1,83739 | 1,58851 | 2 | 7 | 2 | 4 | 9 | 24 |
| **0.5** | 1,93156 | 1,87709 | 1,88744 | 1,83482 | 1,71336 | 1 | 5 | 8 | 6 | 6 | 26 |
| **0.6** | 1,92711 | 1,88486 | 1,89392 | 1,8074 | 1,73788 | 3 | 3 | 7 | 7 | 4 | 24 |
| **0.7** | 1,92281 | 1,88062 | 1,89343 | 1,83975 | 1,72345 | 5 | 4 | 5 | 3 | 5 | 22 |
| **0.8** | **1,92097** | **1,87694** | **1,89516** | **1,84307** | **1,75794** | **6** | **6** | **3** | **1** | **2** | **18** |
| **0.9** | 1,90931 | 1,85789 | 1,89324 | 1,84219 | 1,76077 | 9 | 8 | 6 | 2 | 1 | 26 |
| **1** | 1,9095 | 1,84515 | 1,90011 | 1,83519 | 1,75138 | 8 | 9 | 1 | 5 | 3 | 26 |

**Supp. figure 2**

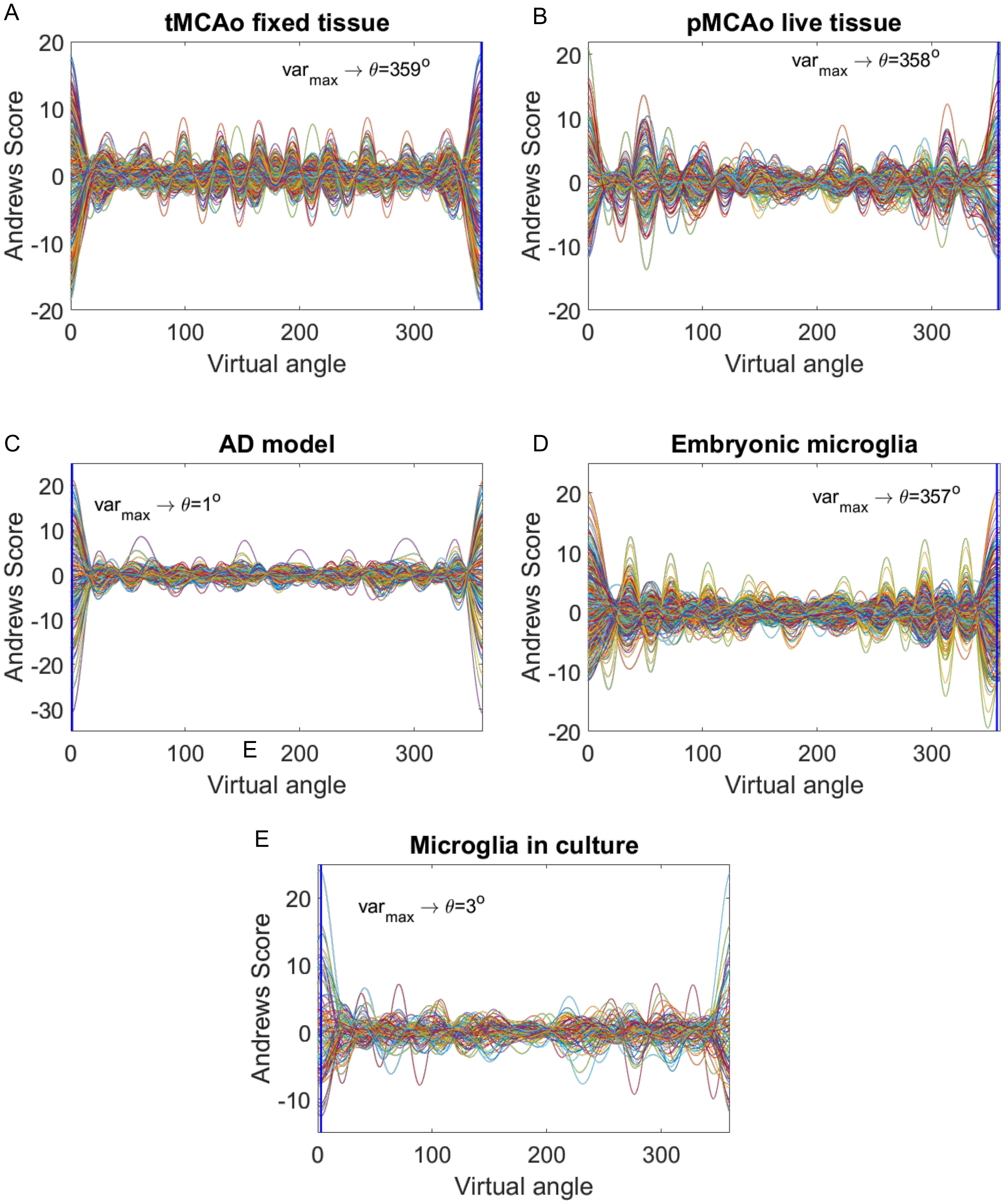

**Supp. figure 3**

### tMCAo fixed tissue

A

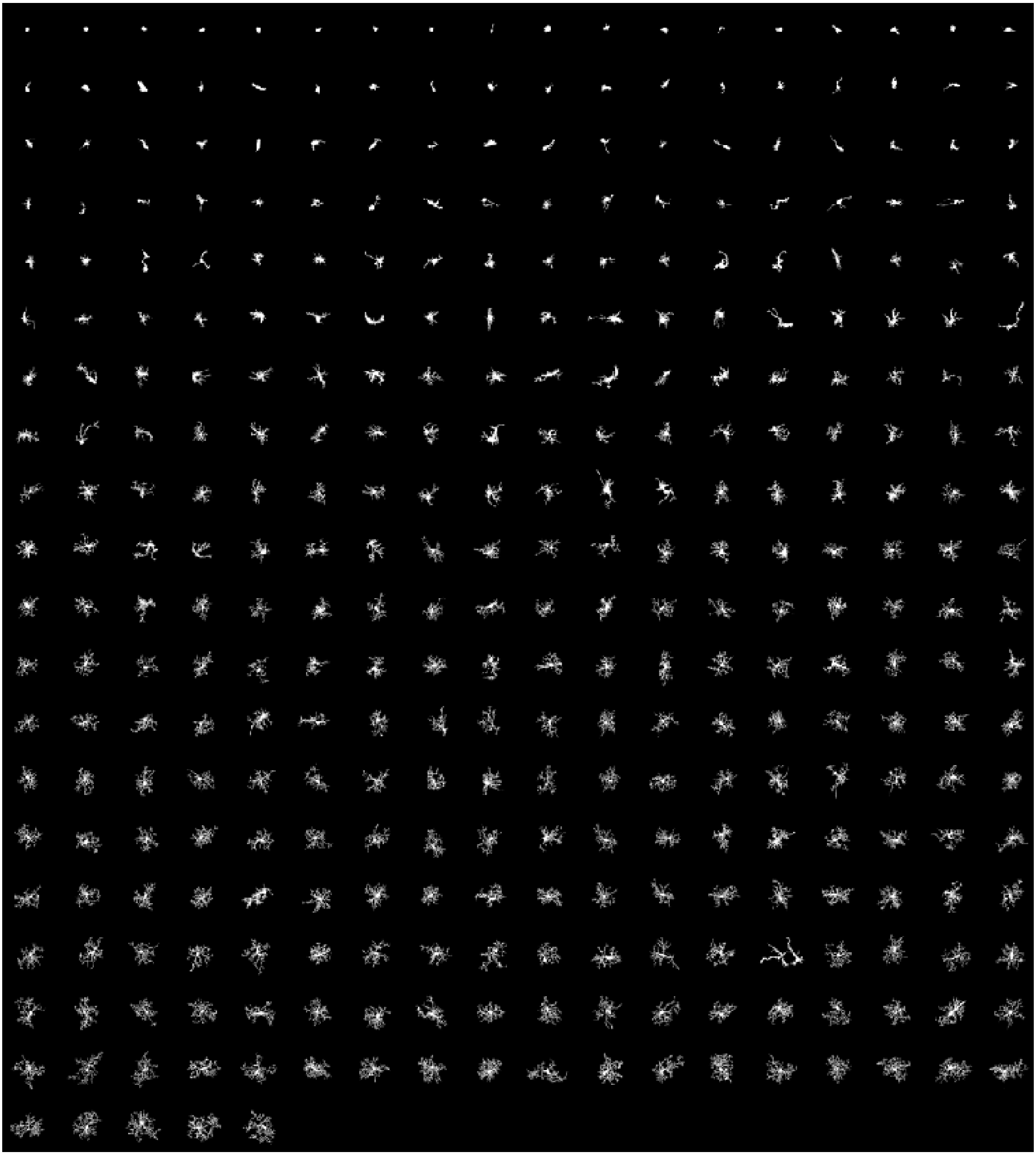

B

**pMCAo live tissue**

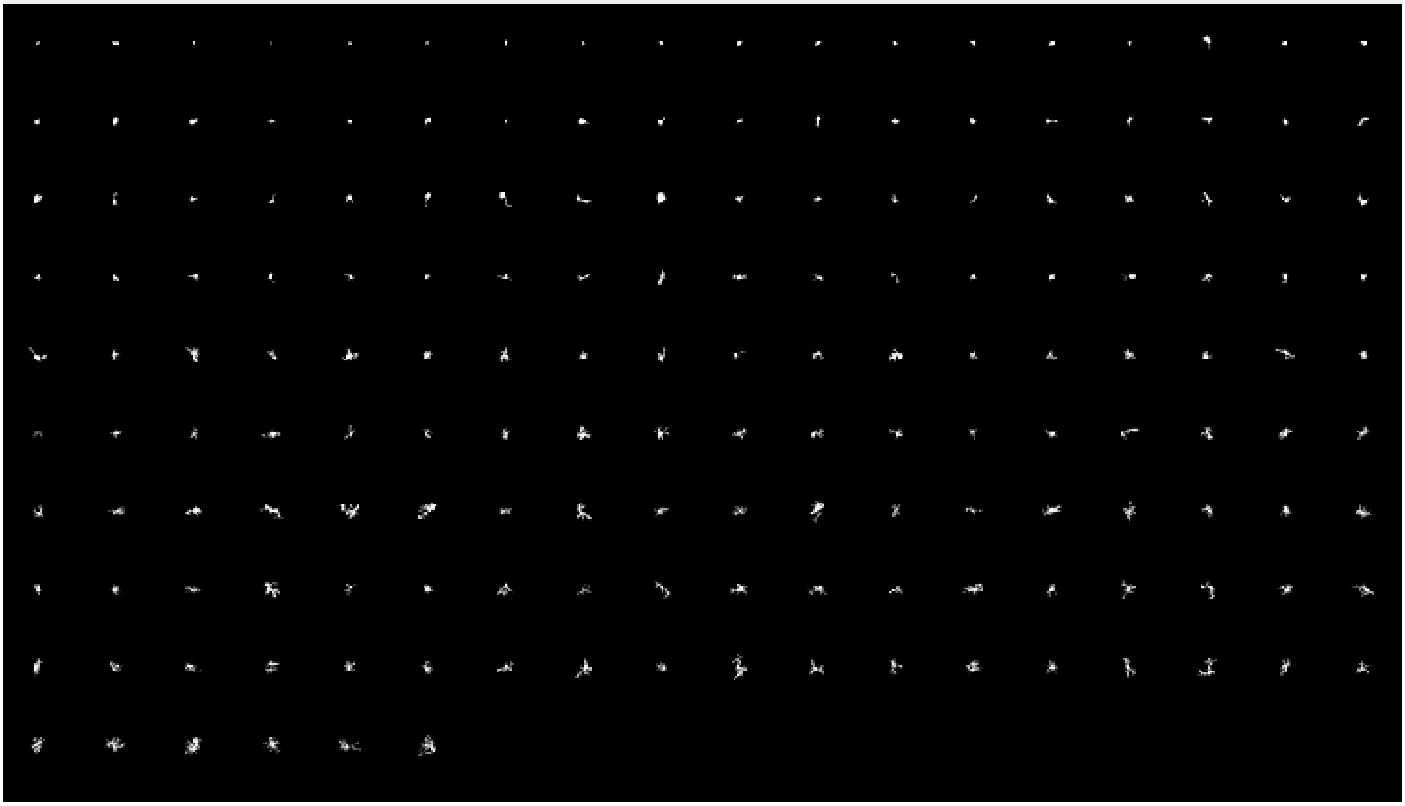

C

**AD model**

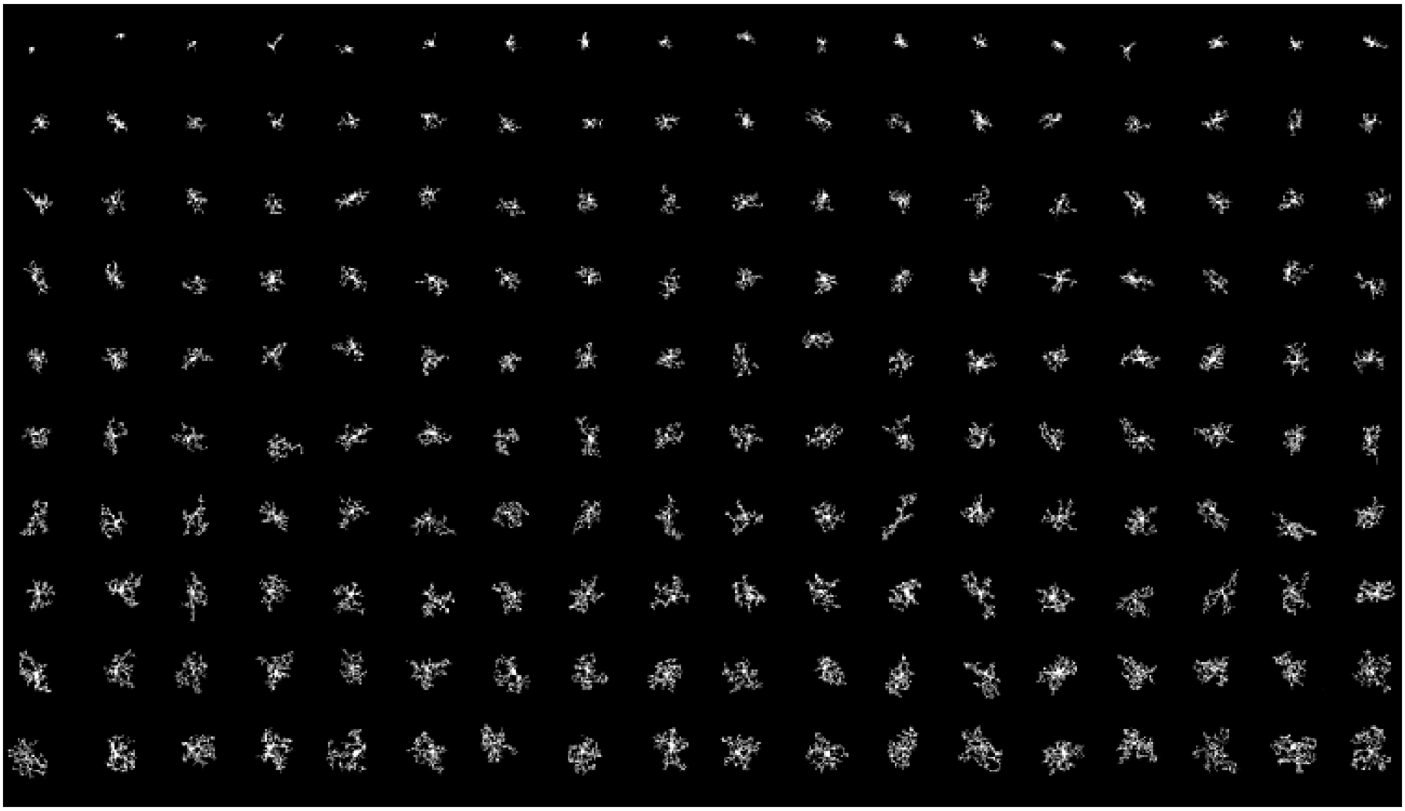

D

**Embryonic microglia**

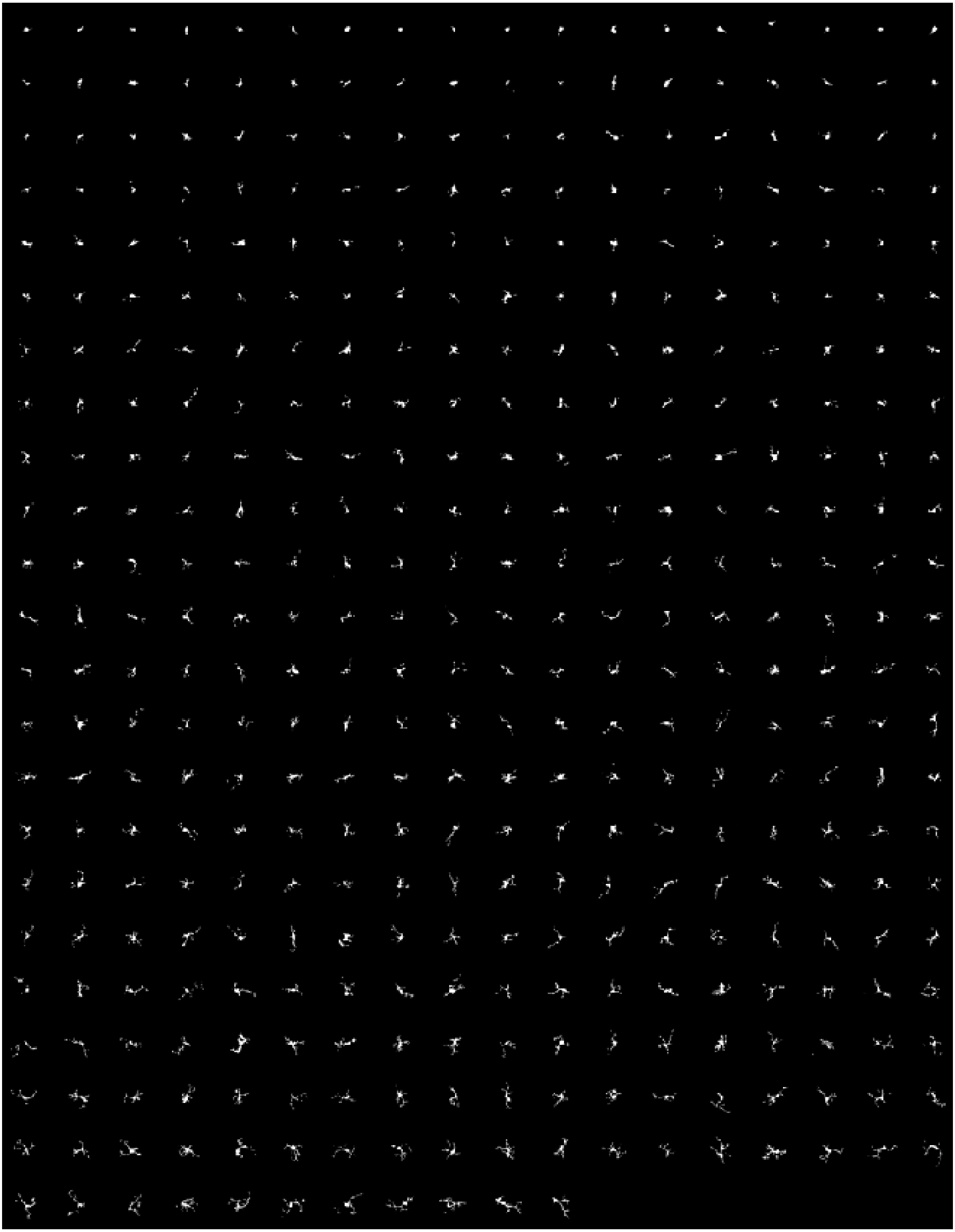

E

### Microglia in culture

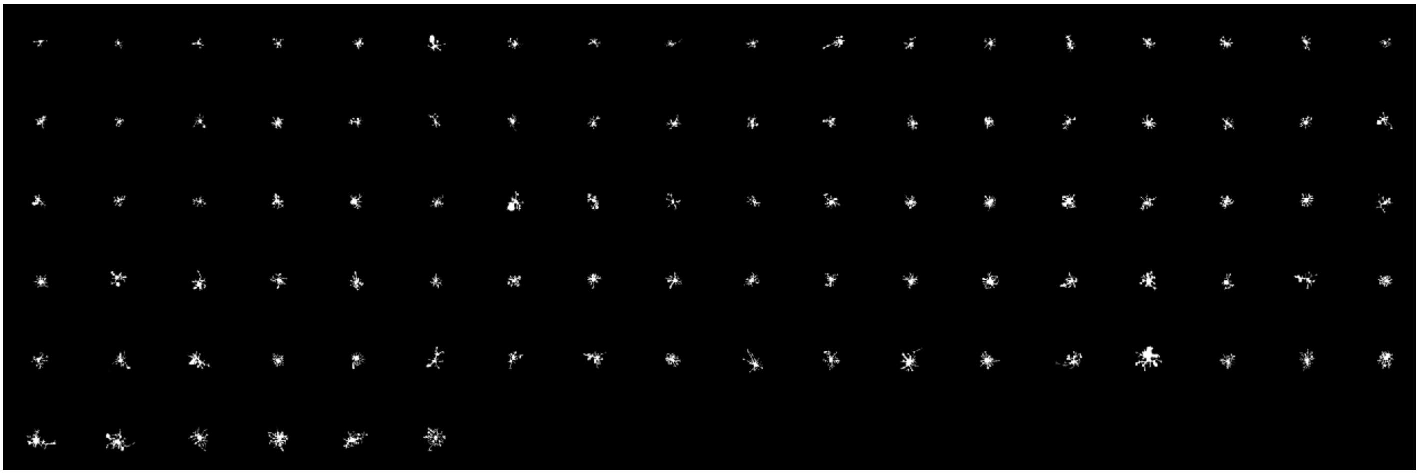

**Supp. figure 4**

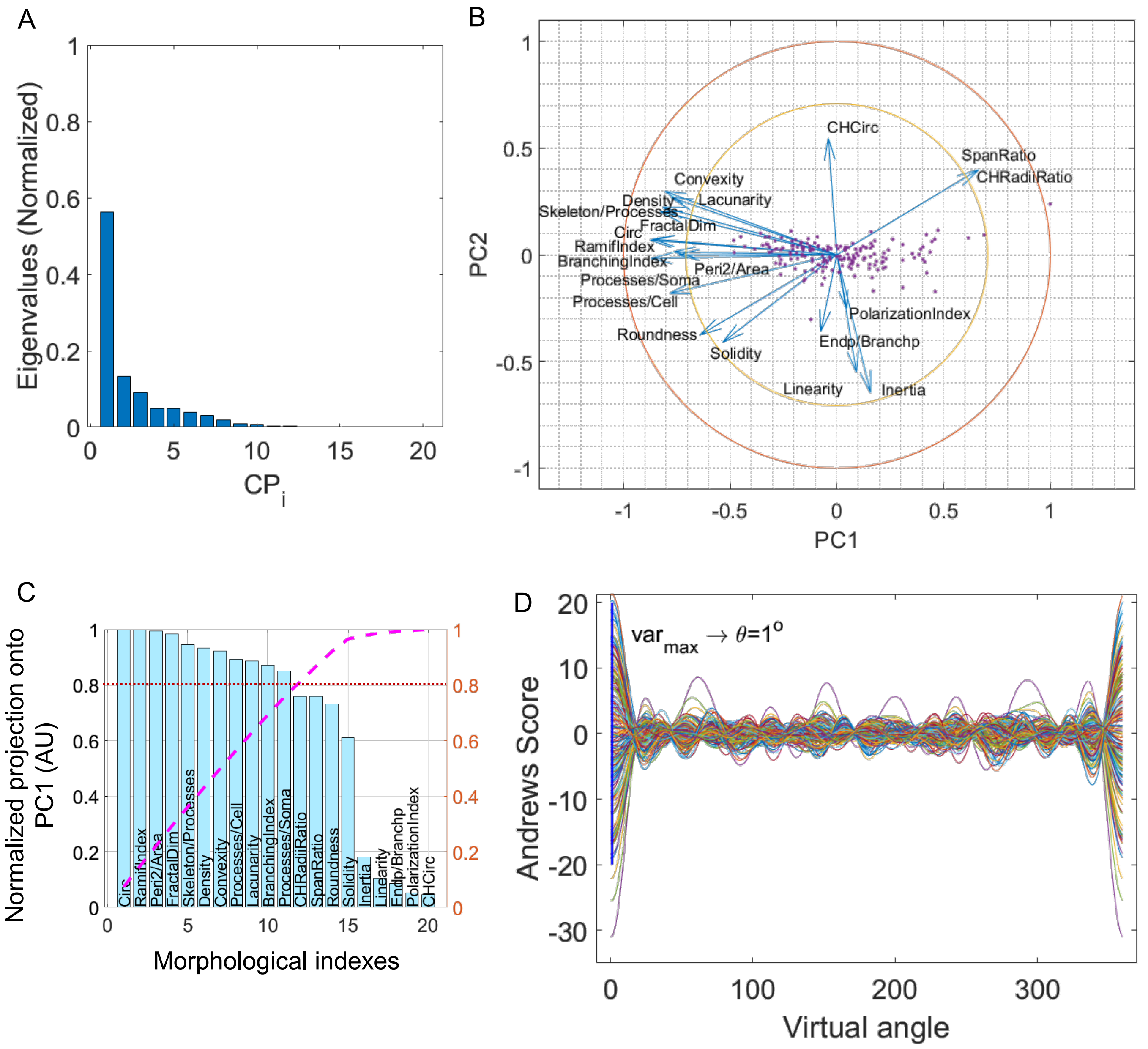

**Supp. figure 5**

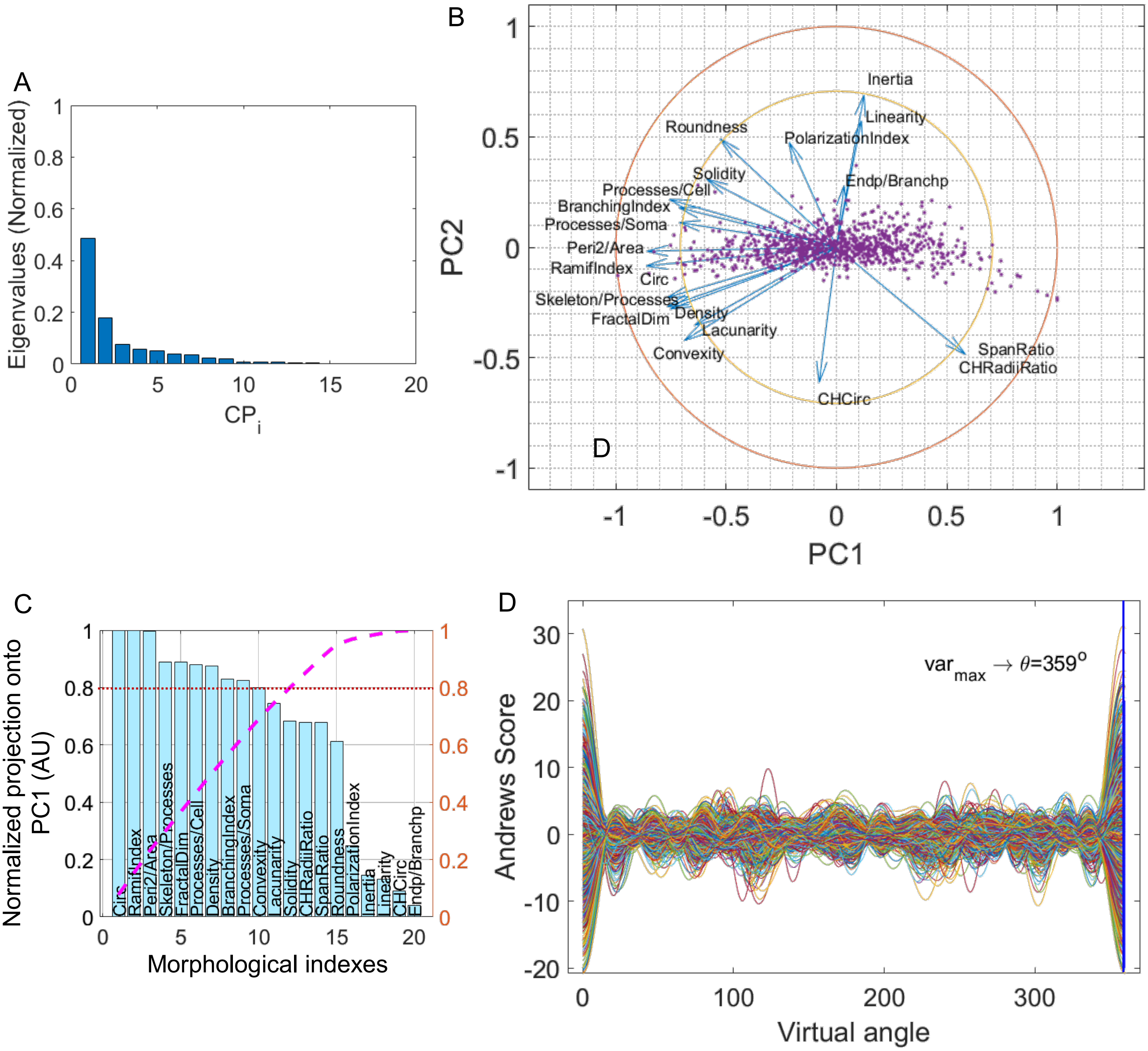

**Supp. figure 6**

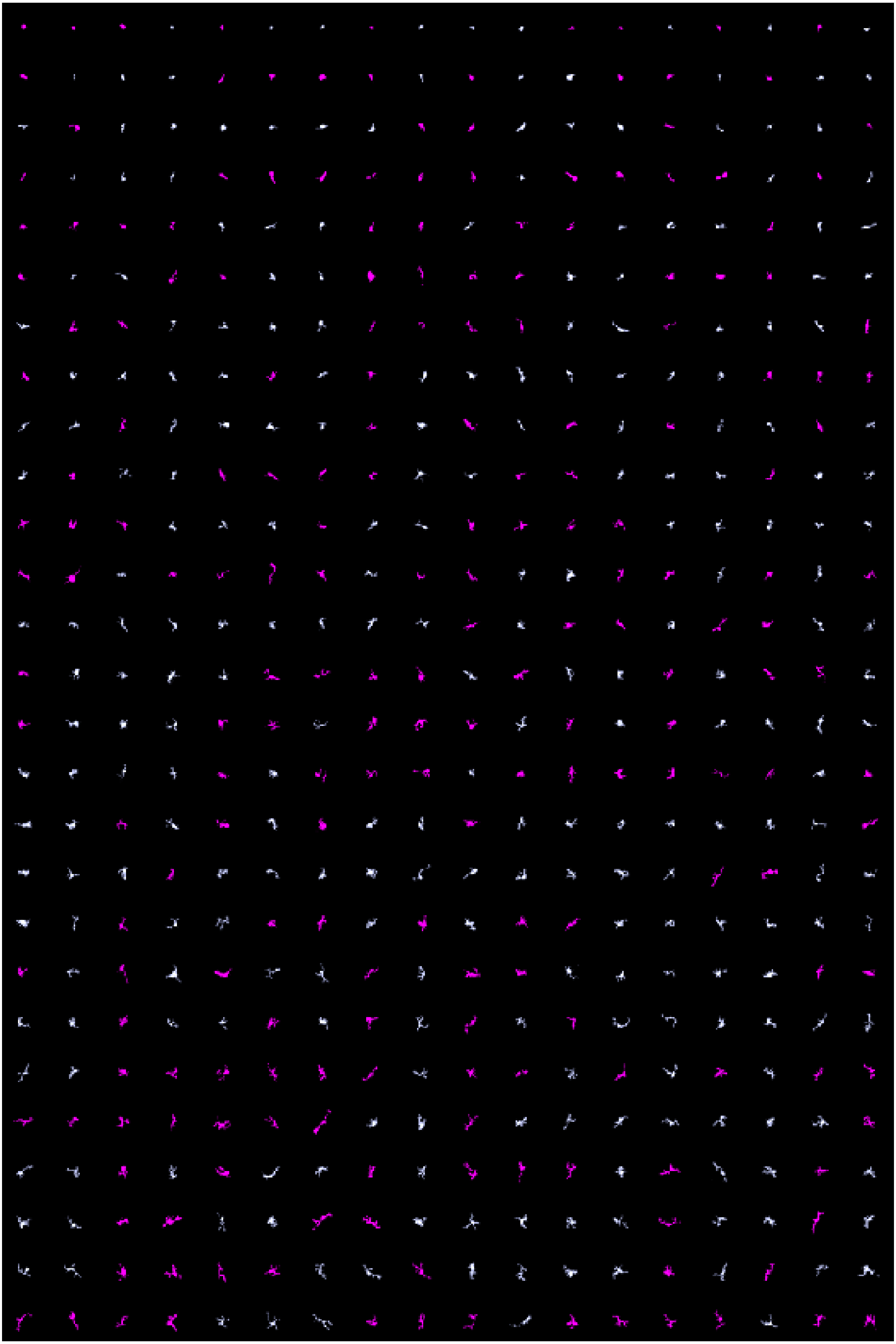

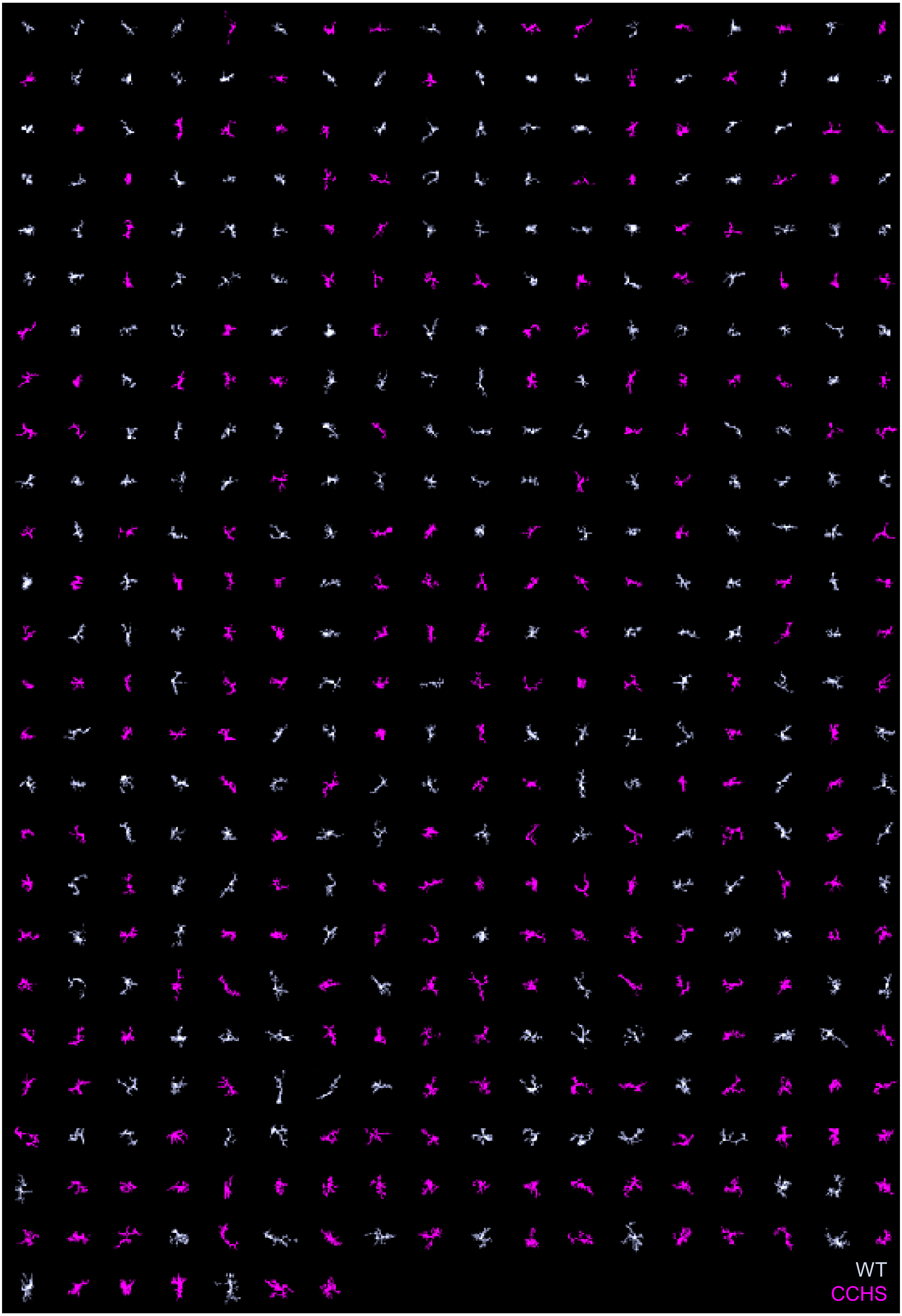

WT
CCHS

Supp. figure 7

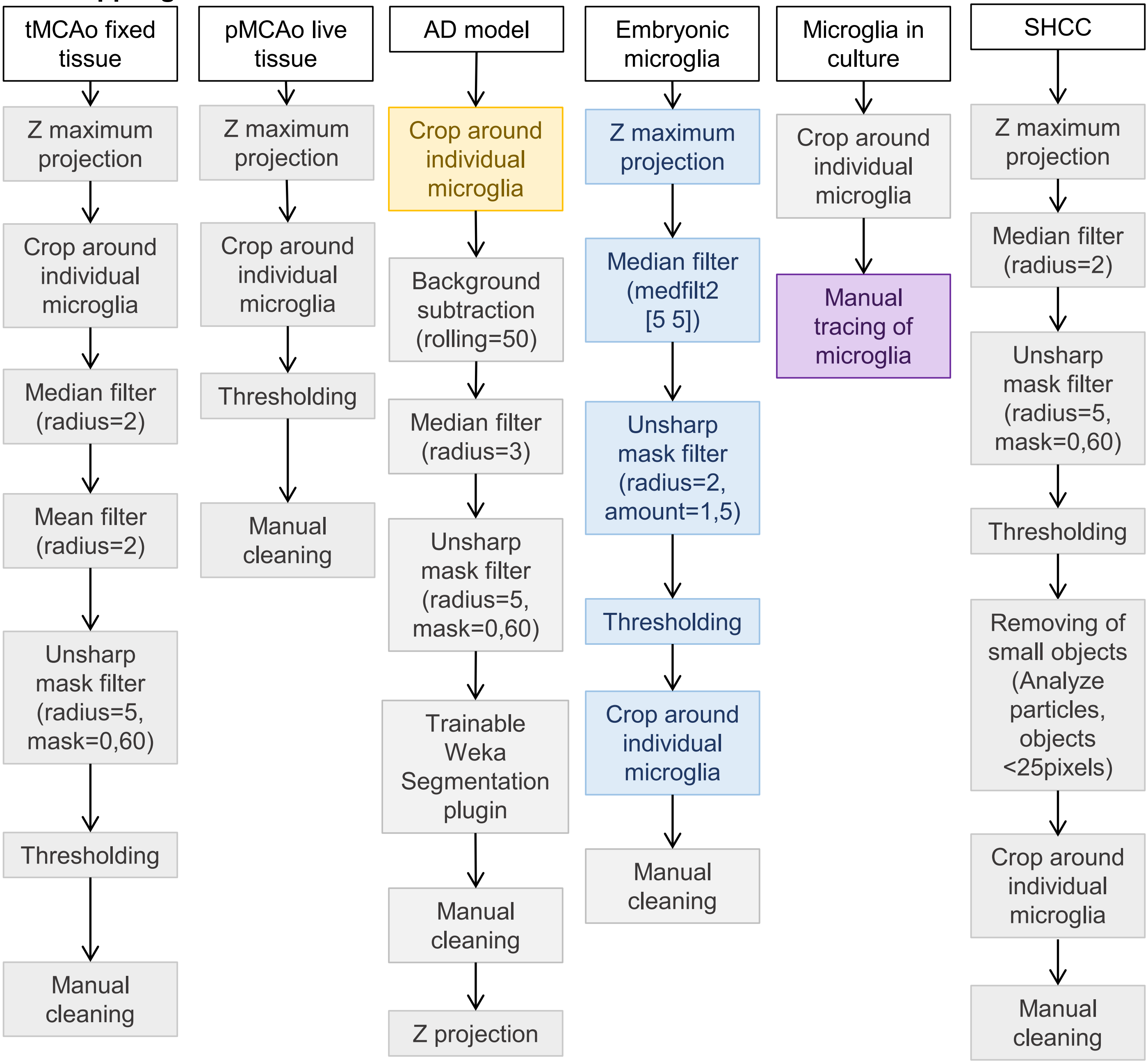

Supp. figure 8

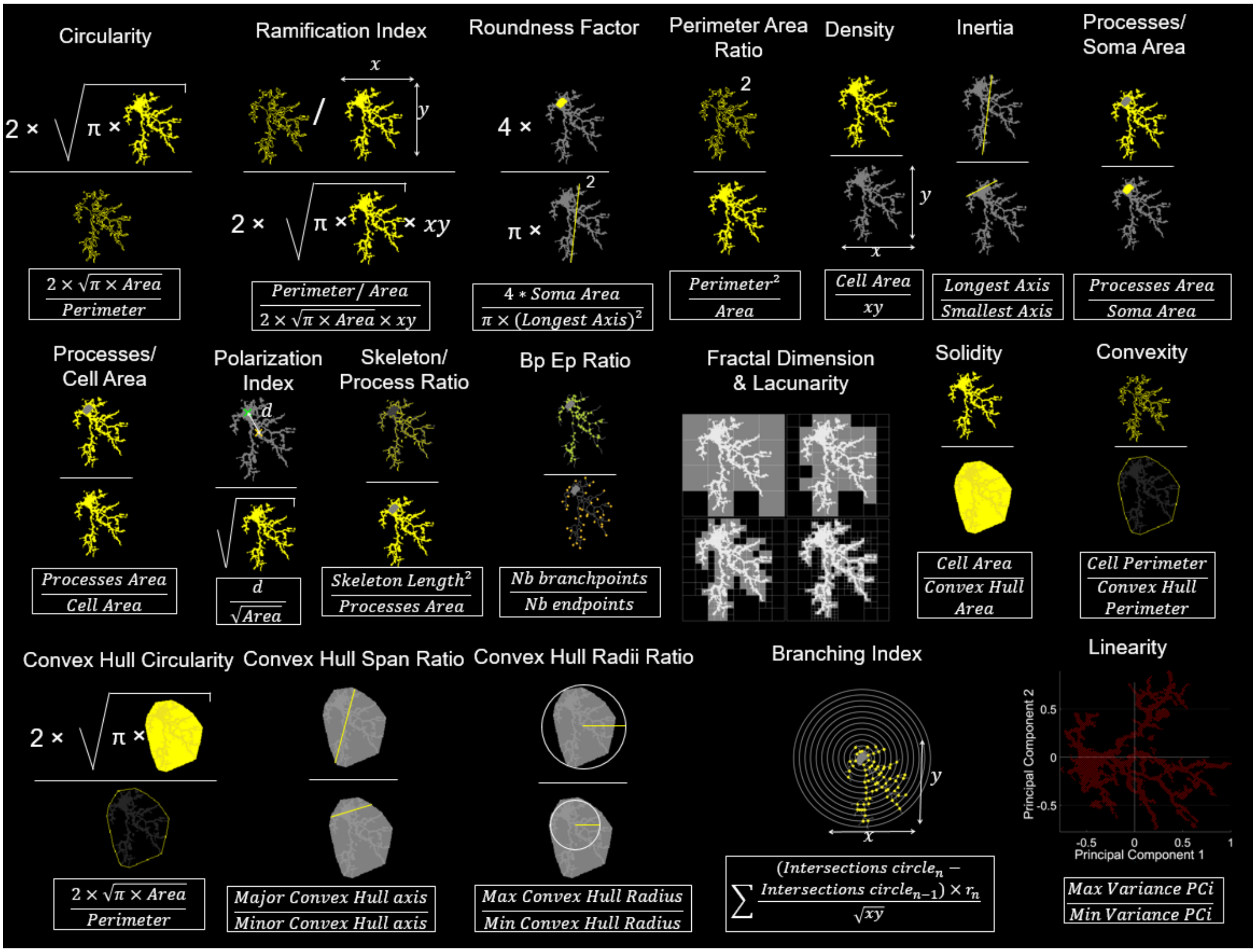
